# Urban larval mosquito surveillance in Antananarivo, Madagascar: spatiotemporal heterogeneity and associated vector dynamics

**DOI:** 10.64898/2026.07.19.739461

**Authors:** Tovo M. Andrianjafy, Johnny A. Uelmen, Lala S. Rafarasoa, Ranto N. Rasolofo, Mamisoa A. Ramahazomanana, Cristina Rafferty, Sarah Zohdy, Ryan M. Carney

## Abstract

Mosquito-borne diseases, such as malaria, remain an important public health problem globally, including on the island of Madagascar. Malaria vector control strategies are typically aimed at adult *Anopheles* mosquitoes in predominantly rural settings. However, very little information is available on larval populations of *Anopheles* and other vector genera, such as *Aedes* and *Culex,* in urban areas, where urbanization creates a multitude of potential larval habitats. Here, the objectives are to identify different types of existing larval habitats and the diversity of vector species in the capital city of Antananarivo, and to study the temporal dynamics and spatial distribution of mosquito vectors of public health interest. From 2022-2023, longitudinal larval surveillance was carried out across all six districts of Antananarivo and consisted of prospecting 2,856 potential larval habitats, of which 653 (23%) were found to contain *Aedes, Culex,* or *Anopheles* larvae. *Anopheles* larvae were reared and morphologically identified using taxonomic keys, and molecular identification by PCR and Sanger sequencing was used for *Anopheles* sibling species determination. A total of 27,418 larval specimens were collected from eight species: *Culex quinquefasciatus* (53%), *Aedes albopictus* (26%), *Cx. pipiens* (14%), *Anopheles gambiae* s.l. (3%, all molecularly identified as *An. arabiensis*), *Cx. univittatus* (1%), *Cx. antennatus* (1%), *Cx. poicilipes* (1%), and *An. coustani* (1%). Geospatial analysis reveals that 55% of larvae came from artificial larval habitats within the city, which harbor lower species diversity than natural larval habitats. Twenty-one different types of larval habitats were observed, and *Cx. quinquefasciatus*, a vector of West Nile virus, was present in all of them. *Aedes albopictus,* vector of arboviral pathogens, prefers artificial habitats such as tires and flower pots, whereas the common rural malaria vector, *An. arabiensis* preferred natural sites like brick pits and rice fields. *An. coustani*, often considered a livestock-associated rural vector, was found to thrive in brick pits, wells, ponds, and in aquatic agriculture. Notably, of the 1,270 *Anopheles* larvae collected, none were found in an artificial container. *Culex quinquefasciatus*, *Cx. pipiens*, and *Ae. albopictus* reached peak abundance in March during the rainy season, while malaria vectors *An. arabiensis* and *An. coustani* showed unexpected abundance peaks during the dry periods, in September and April. Two districts (arrondissements 2 and 5) showed higher abundance and diversity of mosquito vectors than the other districts. The data presented here reveal the spatiotemporal dynamics of malaria and arboviral mosquito vectors in urban Madagascar and highlight the heterogeneity of habitats across the urban ecosystem, with important practical implications for the surveillance and control of mosquito-borne diseases.

## Introduction

Mosquito-borne diseases (MBDs) remain a major public health challenge worldwide, particularly in tropical African countries such as Madagascar, where malaria remains a leading cause of morbidity and mortality (Khezzani et al., 2023; Poungou et al., 2023). In addition to malaria, recurrent outbreaks of dengue and chikungunya occur across the island, placing millions of people at risk and generating substantial social and economic burdens. Effective prevention and control of MBDs depend on robust vector surveillance systems that identify when, where, and which mosquito vectors are present. Historically, surveillance and control efforts targeting indoor-biting and indoor-resting *Anopheles* mosquitoes in predominantly rural settings, through interventions such as indoor residual spraying (IRS) and insecticide-treated bed nets (ITNs), have contributed substantially to reductions in malaria burden (Bhatt et al., 2015; WHO, 2024). However, as urbanization accelerates and the risk of multiple MBDs increases, there is growing recognition of the need to better understand mosquito vector communities in urban environments. Larval habitats of *Anopheles* mosquitoes may overlap with those of other medically important vectors, including *Aedes* species that transmit dengue and chikungunya, and *Culex* species that transmit West Nile virus, creating critical opportunities for integrated urban vector surveillance and management. Despite this potential, knowledge of mosquito species diversity, spatiotemporal dynamics, and larval habitat characteristics in urban areas of Madagascar remains limited.

Urban environments are full of anthropogenic activities, such as construction and agriculture, that generate stagnant water, which can be favorable to the development of mosquito juveniles. One key feature of urban areas is also high human population density, which corresponds to a high host blood meal availability and a high risk of transmission of certain vector-borne diseases. This is the case in Antananarivo, the capital of Madagascar, which is home to around 4 million people with a 4.56% growth rate (UN World Urbanization Prospects, 2024). In recent years, it has been marked by significant urban expansion. The potential interaction between humans and mosquitoes in Antananarivo is of potential public health concern. Urbanization can profoundly modify ecosystems by transforming natural habitats into artificial environments, subsequently influencing the distribution of larval habitats and promoting vector proliferation (Wilke *et al*., 2019). Dengue and chikungunya epidemics have occurred in many towns throughout Madagascar, such as urban transmission of these viruses in Mahajanga and Toamasina (Rabarijaona *et al*., 2006; Ratsitorahina *et al*., 2008; Randrianasolo *et al*., 2010). In recent years, the landscape of malaria, previously thought to be a predominantly rural disease in Africa, has shifted to include urban areas (WHO urban malaria framework, 2022; Sinka *et al*., 2020; Hamlet *et al*., 2022). In Madagascar, both malaria cases and deaths doubled from 2019 to 2020, to their highest numbers in over two decades (WHO 2021). Madagascar remains among the 20 countries with the highest rates of malaria with respect to estimated cases (2,861,319) and deaths (395) in 2024 (WHO, 2024).

*Anopheles stephensi*, a competent vector of malaria in urban areas endemic to Asia and the Arabian Peninsula, has become successfully established as an invasive species in African countries, including Djibouti, Ethiopia, Sudan, Nigeria, Eritrea, Ghana, Kenya, and Somalia (WHO, 2023; Ochomo *et al*., 2023; Ali *et al*., 2022; Ahmed et al., 2021; Carter et al., 2021; NIMR, 2022). This species is now of great concern due to the sharp increase in urban malaria incidence associated with the presence of invasive *An. stephensi* in Djibouti, and the evidence of the vector’s transmission of malaria parasites in urban Ethiopia (De Santi *et al*., 2019; Hamlet et al., 2022; Emiru *et al*., 2023). To address the threat of *An. stephensi*, enhanced larval and adult mosquito surveillance in all African countries has become a priority for malaria programs for early detection and rapid response (WHO vector alert 2023 update, *PMI* 2023). This monitoring is not limited to rural areas but extends to peri-urban and urban areas, where *An. stephensi* thrives, especially in large cities like Antananarivo, for early detection and rapid response to *Anopheles stephensi* and to better understand urban malaria dynamics (WHO urban malaria framework, 2022).

The main objective of this work is to conduct enhanced longitudinal larval mosquito surveillance in Antananarivo, Madagascar, to determine the 1) diversity and abundance of *Anopheles* and other vectors and associated larval habitats, 2) spatiotemporal heterogeneity of mosquito species, and 3) impacts of socioenvironmental factors such as population density, precipitation, land use/land cover, temperature, socioeconomic, and built environment indicators on larval habitats and species composition. These data can be used to better understand the spatial risk of MBD transmission in this major urban center and to support initial surveillance to determine if *An. stephensi* is in Madagascar (Carney *et al*., 2023), a location considered to be at potential risk of *An. stephensi* introduction (Ahn *et al*., 2023).

## Materials and methods

### Study site

The study was conducted in Antananarivo, Madagascar’s capital. The city is divided into six arrondissements (districts; **Figure 1A**) and 192 fokontany (villages). It is located at an altitude of 1,280 m in the high plateau region and has an area of 107 km^2^. The population exceeds 3 million inhabitants with a density of approximately 30,000 inhabitants per km^2^. Antananarivo has a tropical altitude climate. It is dotted with rice fields, ponds, lakes, rivers, and abandoned artificial containers scattered across the city. All habitats were included because they are potentially important larval habitats for mosquito vectors.

**Figure 1.**
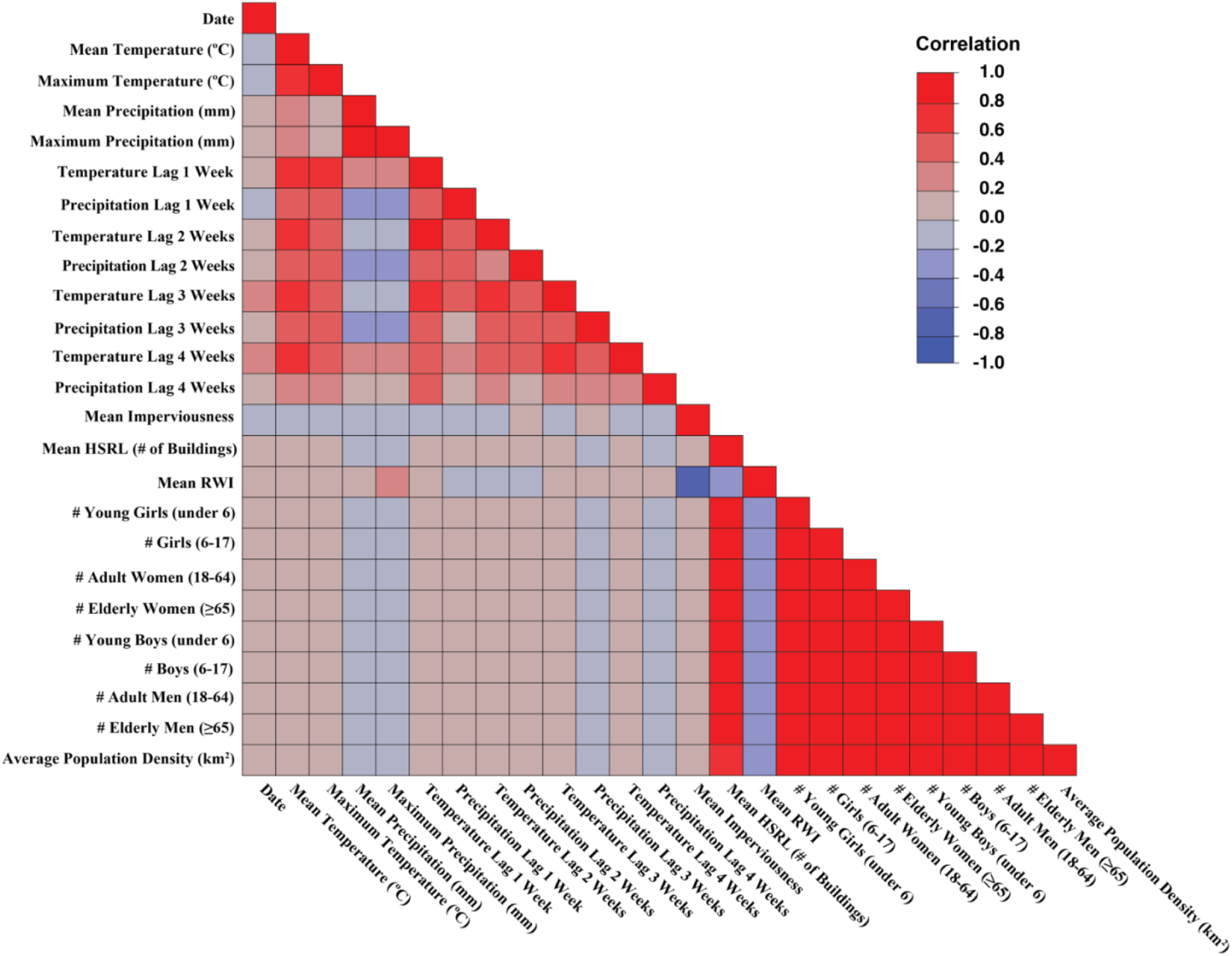
Correlation plot of predictors assessed for model inclusion. The following eight population characteristic variables were removed from further assessments: # Young Girls (under 6), # Girls (6-17), # Adult Women (18-64), # Elderly Women (65+),

### Larval habitat surveillance

Prospecting for larval habitats was conducted across all six arrondissements weekly for 12 months (July 2022 to July 2023). Surveillance consisted of carefully inspecting all potential larval habitats, both artificial and natural, for Culicidae larvae. Sites were selected after observing different locations in each arrondissement on Google Maps. 20 to 30 houses within the arrondissement were randomly selected for surveys (with the proprietor’s permission; otherwise, we selected another house in the same arrondissement). The parameters observed during the survey were: presence or absence of larvae or pupae, larval habitat categorization (natural or artificial), type or specific descriptive characteristics of the larval habitats (container, tire, rice field, etc.), distance to human dwellings (close <30m or far ∼30-200m), and geographical coordinates of each habitat inspected. Habitats considered natural include: bamboo, brick pit, coconut waste, drainage channel, footprint, pond, puddle, rice field, tree trunk, water basin, water well, and watercress. Artificial habitats include: barrel, cut plastic canister, flower pot, metal container, plastic bowl, plastic bucket, small plastic container, tire, and used bottles.

### Mosquito collection and identification

Mosquito larvae and pupae were collected directly from the larval habitats using pipettes, ladles (350ml), and white plates (500ml). Our systematic search of all potential larval habitats in the inspected areas aimed to inventory the species present and report their abundance as the cumulative number of individuals. While not the focus of the study, larval density was also estimated. In natural habitats, 20 dips using a 350mL ladle were conducted to estimate larvae/L. For artificial habitats >1 L, two 350mL dips were conducted to estimate larvae/L. For artificial habitats < 1 L, all larvae were counted and the volume estimated to obtain a number of larvae/L. For example, the number of larvae from 0.5 L of water was multiplied by 2. In large natural habitats, dips were conducted along the four sides and the center of rectangular habitats to ensure representative sampling of the habitat. All specimens were placed in labeled containers with habitat water and placed in a cooler before being brought back to the laboratory for identification and rearing to adulthood.

### Mosquito larval rearing to adulthood

Mosquito rearing was conducted at the insectary under the following conditions: 25 ± 3°C, 70% relative humidity, and a photoperiod of 14/10 hours. For each mosquito genus, larvae were transferred to separate tanks (32 × 32 × 12 cm^3^) containing larval habitat water. They were fed a vitamin-rich powder (Steach® fish food), a pinch of which was added to each rearing tank every three days. Larvae were put into a tank for each larval habitat surveyed. The pupae were separated from the larvae and placed in cups containing breeding water. Then, they were placed inside a Gauze cage (35 × 35 × 35 cm^3^) while waiting for their emergence. Adult females and males were kept together and fed with cotton soaked in a 6% sucrose solution suspended inside the cage.

### Morphological species identification

The larvae were first sorted by genus, and then the 4^th^ instar was morphologically identified under a binocular magnifier (AmScope®) down to the species level using the identification keys (Robert *et al*., 2022), where possible. When the adults emerged, re-identification based on the morphological characteristics of the individuals was carried out to confirm the previous identification results during the larval stages, this time using keys intended for adults (Grjebine, 1966; Coetzee, 2021).

### Molecular species identification by PCR and Sanger sequencing

Only sibling species within the *Anopheles gambiae* complex were identified to the molecular species level using PCR (Wilkins *et al*., 2006). From our urban surveys, 10 larvae and 162 adults (161 reared from larvae) were molecularly processed using PCR and Sanger sequencing for species confirmation. The PCR method used to amplify the ITS2 region of the *An. The gambiae* complex was adapted from the Wilkins et al. protocol as follows: 1 μl DNA, 25 μM of each primer IMP-UN, AR-3T, GA-3T, ME-3T, IMP-S1, IMP-M1, QD-3T, and 2X Accustart® PCR MasterMix (Quantabio) in an 11.5 μl reaction. PCR conditions were the following: 94°C-3 min; 30 cycles of 94°C-30 sec, 60°C-30 sec, 72°C-30 sec; 72°C-5 min. Products were visualized on a 1.5% GelRed-stained agarose gel, run at 100V for approximately 60 min.

### Amplification and sequencing

Molecular analysis was performed on the samples by amplifying the ITS2 gene using the primers and conditions described originally by Beebe *et al*. (1995) and adapted as follows: 10μM of each primer, 2X Accustart®PCR MasterMix (Quantabio, Beverly, MA), and water for a total reaction volume of 12 μl (Beebe & Saul, 1995). Temperature cycling was as follows: 95°C-5 min; 35 cycles of 95°C-30 sec, 56°C-45sec, 72°C-1 min; 72°C-10 min. Three microliters of PCR product were run on 2% GelRed-stained agarose gel, run at 90V for 60 min to confirm successful PCR assays. PCR products were then cleaned using ExoSAP and sequenced using Sanger technology with an ABI BigDye TM Terminator v1.1 cycle sequencing kit (Life Technologies, Santa Clara, CA) according to the manufacturer’s recommendations, and the resulting sequences were run on an ABI 3500xL Genetic Analyzer. Sequences were cleaned and analyzed using SeqManPro v.17.3.0 (DNAstar Lasergene, Madison, WI). ITS2 sequences from the samples were submitted as queries to the National Center for Biotechnology Information’s (NCBI) web-based Basic Local Alignment Search Tool (BLAST) against the nucleotide collection in Genbank under default parameters (max High scoring Segment Pairs (HSP). The top sequences from NCBI that formed HSPs with the queries were identified as matches.

### Socioenvironmental factors and data sources

All external environmental, human demographic, and weather data are freely available for public use and were extracted for each of the six arrondissements. Weather data were acquired from the National Science Foundation (NSF) National Center for Atmospheric Research (NCAR) Global Surface Observational Weather Dataset (NCEP 2004). Daily minimum and mean temperature (℃) and mean precipitation (mm) were acquired between June 2022 and July 2023 from each of the two weather stations in Antananarivo. To provide continuous coverage across our study area, all daily weather pixel values were interpolated (using the inverse distance weighted (IDW) method) in ArcGIS Pro (version 3.2, ESRI, Redlands, CA USA). Estimated daily weather values were then converted to weekly temperature lags (0-4 weeks) with respect to the collection date for each larval observation. Land use and land cover (LULC) data were acquired from the Copernicus Global Land Service (Copernicus 2019). Data, represented as the average pixel value for each 100-meter-resolution grid in 2019, were extracted for all arrondissement areas. Human socioeconomic data, consisting of population by gender and age and a socioeconomic index of relative wealth (RWI), were acquired from the Humanitarian Data Exchange (HDX 2020). Human population data consisted of high-resolution (100 m) geographic-embedded tiff images (geotiff) of gender and age group (children >5, youth (15-24), men and women (>18, including women of reproductive age (15-49)), and elderly (adults >60)) by pixel (v1.75.1 PY3), for the year 2019. Relative wealth was available as a comma-separated value (CSV) dataset with corresponding latitude and longitude coordinates. Similarly to the previously described analysis with weather data, all RWI point values (n = 43,640) were converted to a continuous raster layer by interpolation (IDW) in ArcGIS Pro. Additionally, to approximate the footprint and environmental impacts of human development, we extracted datasets on human population density (km^2^), imperviousness, and the human-built environment for each of the six arrondissements. Human population density (all ages and genders combined) was obtained from the WorldPop Hub as 1 km-resolution geotiff data for 2020 (WorldPop 2020). Global gridded man-made imperviousness and human-built environment data (both at 30 m resolution) were acquired from the National Oceanic and Atmospheric Administration (NOAA) Socioeconomic Data and Applications Center (SEDAC) dataset (SEDAC 2010).

### Statistical Analyses

Mosquito collections were reported as a singular point in space and time. Given mosquito biology, behavior, and physiology, we created a 1 km buffer around each collection location. This 1 km buffer is a reasonable estimate of the average female flight range across the species evaluated in this study, while accounting for differences between genera. Within this buffered zone, we overlaid, extracted, and merged all aggregated raster data (continuous) with categorical data to account for the heterogeneity of predictor variables within the proximity of each trap location.

Before any analyses, we evaluated collinearity among variables, leading to the removal of eight predictors from further analysis (**Figure 1**). In total, our final dataset comprised 24 independent variables of interest pertaining to the dependent variable, the number of collected larvae (**Table S1**). A descriptive analysis evaluated each predictor by outcome (mosquitoes collected by species), providing mean and sum (categorical variables only) values over the 1-year study period. Univariate analyses were conducted to evaluate the overall relationship between each predictor and outcome combination (assessed as the portion of covariance inclusion, e.g., contribution). Models for each dependent variable (eight mosquito species + pooled) were evaluated using the Bootstrap Forest method (169 layers, 7 splits per tree, 653 training rows) and reported as best-fit based on linear correlations of predicted vs. actual points (r^2^). Within each best-fit model output, the amount of variance (sum of squares) captured by each predictor was compared and contrasted across models. Spatial autocorrelation for all mosquito species was assessed using Moran’s I. Kernel density distributions for all collected mosquitoes by species show the geographic relationship in abundance, categorized by quintiles and color-coded (spectrum from hot/frequent (red) to cold/few (blue)). All statistical analyses were generated using JMP Pro (17.0.0) and SAS (9.4) software (SAS Institute, Inc., Cary, NC, USA).

## Results

The total number of inspected potential larval habitats in Antananarivo across the year was 2,856, of which 23% (n=653) had larvae and 77% (n=2203) did not.

### Mosquito diversity and abundance

A total of 27,742 specimens of Culicidae larvae were collected during the 12 months of monitoring in the city of Antananarivo. These larvae belonged to three genera: *Aedes*, *Anopheles*, and *Culex*, of which eight species were morphologically and molecularly identified: *Cx. quinquefasciatus* was the most abundant (53%), followed by *Ae. albopictus* (26%), *Cx. pipiens* (14%), *An. arabiensis* (3%), *Cx. univittatus* (1%), *Cx. antennatus* (1%), *Cx. poicilipes* (1%) and *An. coustani* (1%). One adult mosquito was collected using a mouth aspirator in a kitchen near a cowshed and was suspected to be *An. stephensi* based on wing patterns, but was molecularly identified to be *An. arabiensis*. Two larvae reared to adults were morphologically identified as *An. coustani*, which were molecularly confirmed by ITS2 Sanger sequencing. 159 larvae reared to adults were morphologically identified as *An.* gambiae s.l.; 157 were molecularly confirmed as *An. arabiensis* by ITS2 Sanger sequencing and *An. gambiae* complex by species ID PCR, and two specimens had no DNA sequence match and failed to amplify. 10 larvae identified as *An. gambiae* s.l. failed to amplify.

In general, 55% (n=15,199) and 45% (n=12,543) of the larvae collected came from natural and artificial habitats, respectively. Regarding each species, most of them were found in natural habitats, except for *Cx. antennatus* and *Ae. albopictus* which were predominantly found in artificial habitats **(Table S1).** The percentage of larvae collected from permanent larval habitats was 56% (n=15,444), and the remaining 44% (n = 12,298) were from temporary larval habitats. Most species preferred permanent larval habitats over temporary larval habitats, except for *Cx. antennatus* and *Ae. albopictus*. The abundance of larvae varied by locality, with a larger number of individuals collected farther from dwellings (54%) than close to dwellings (46%). However, unlike the other species, *Ae. albopictus* was predominantly found near human habitation.

Tires (n = 3,779 individuals per habitat) and flower pots (n = 3,410 individuals per habitat) had the highest numbers of larvae among the sites surveyed, followed by watercress habitats, puddles, drainage channels, and rice fields (between 2,000 and 3,000 total individuals collected per habitat). The sites with the fewest larvae were tree trunks, footprints, metal containers, used bottles, and coconut waste, each with fewer than 200 individuals. *Cx. quinquefasciatus* was present in all larval habitats but was most abundant in watercress and puddles (>2,000 individuals), whereas it was relatively rare in other larval habitats, such as footprints, metal containers, plastic bowls, and tree trunks. *Cx. pipiens* was abundant in drainage channels, puddles, watercress, ponds, rice fields, but absent in some habitats, including bamboo, plastic buckets, metal containers, and used bottles. *Cx. poicilipes* and *Cx. univittatus* were collected only in rice fields and brick pits with low numbers (< 200 individuals). *Cx. antennatus* is similar but also occurs in other larval habitats, such as tires and cut plastic canisters. *Anopheles arabiensis* was collected the most in brick pits (n = 404) and rice fields (n = 211). This malaria vector species was also present in water basins, water storage wells, watercress (50-100 individuals), ponds, puddles, and footprints (<50 individuals). *Anopheles coustani* was found in brick construction pits, ponds, rice fields, water basins, water wells, and watercress. *Ae. albopictus* was found in high abundance in flower pots and tires (>2,000 individuals), followed by plastic buckets, bamboo, and cut plastic canisters (around 500 individuals). This species was also found on some tree trunks, in small plastic containers, and in barrels (**Figure 2**). In terms of diversity, natural larval habitats such as rice fields, puddles, and watercress harbor more species (8) compared to artificial larval habitats with only two or three species.

**Figure 2.**
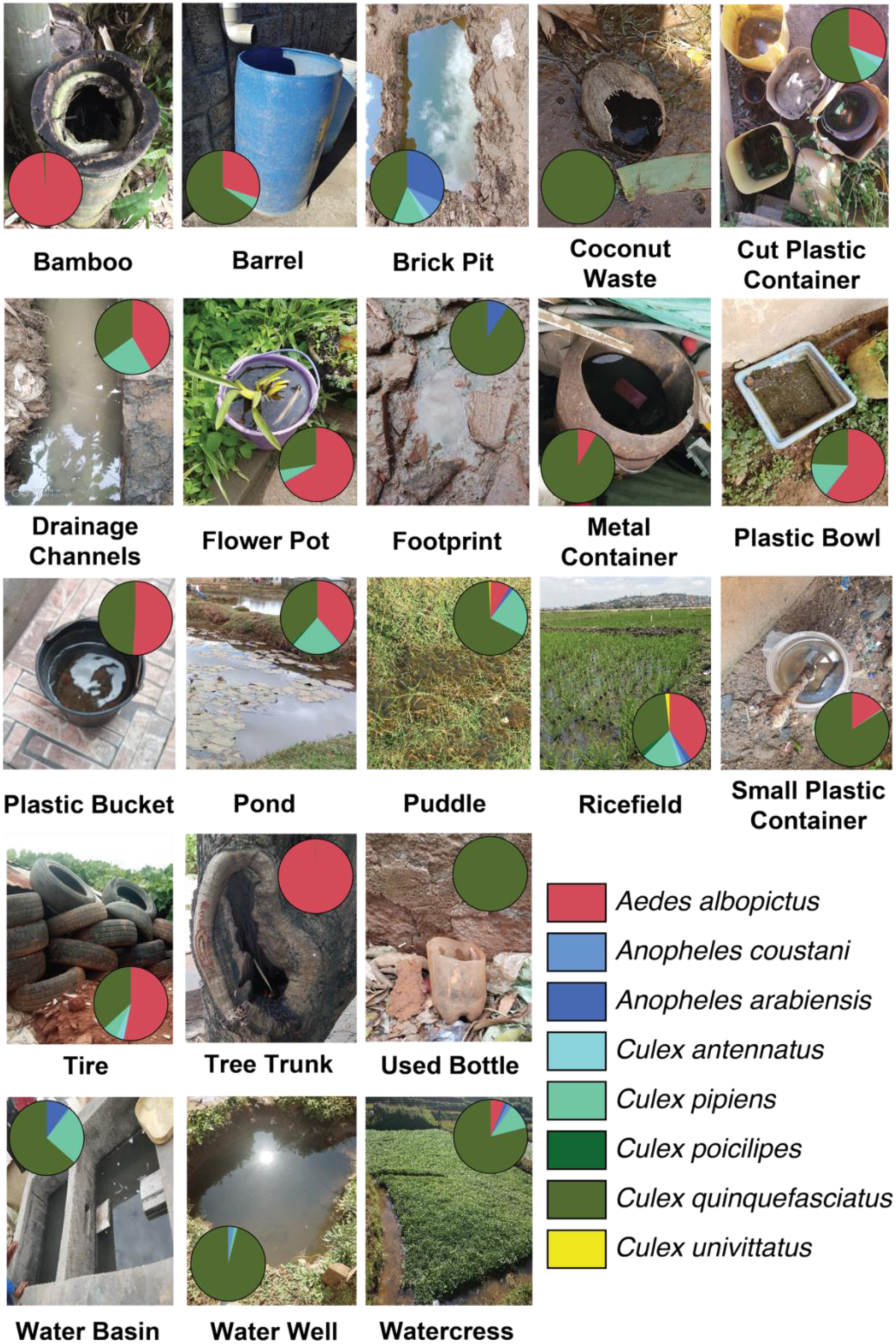
Mosquito vectors were collected across a wide range of natural and artificial larval habitats in Antananarivo, Madagascar, from July 2022 to July 2023. Some habitats, such as brick pits, rice fields, and ponds, contained a higher diversity of species, while others, such as used bottles and coconut waste, only contained one species (*Cx. quinquefasciatus*).

When larval counts are expressed as larval density (larvae/L) rather than raw abundance, the most productive larval habitats shift toward small, artificial containers. *Ae. albopictus* and *Cx. quinquefasciatus* reached their highest larval densities in containers such as tires, small plastic containers, buckets, and flower pots, whereas expansive natural habitats—rice fields, ponds, and watercress—yielded high total abundance but comparatively low larval density owing to their much larger water volumes (Figure 3, Figure 4). This contrast was most pronounced for the malaria vectors *An. arabiensis* and *An. coustani*: Although collected in appreciable numbers from rice fields (n = 211) and brick pits (n = 404), their larval densities remained low, consistent with dilution across large water bodies. Because raw abundance can overstate the importance of a single large habitat (e.g., a single rice field or pond), larval density more directly highlights the container habitats where source reduction and larviciding are likely to be most impactful.

**Figure 3.**
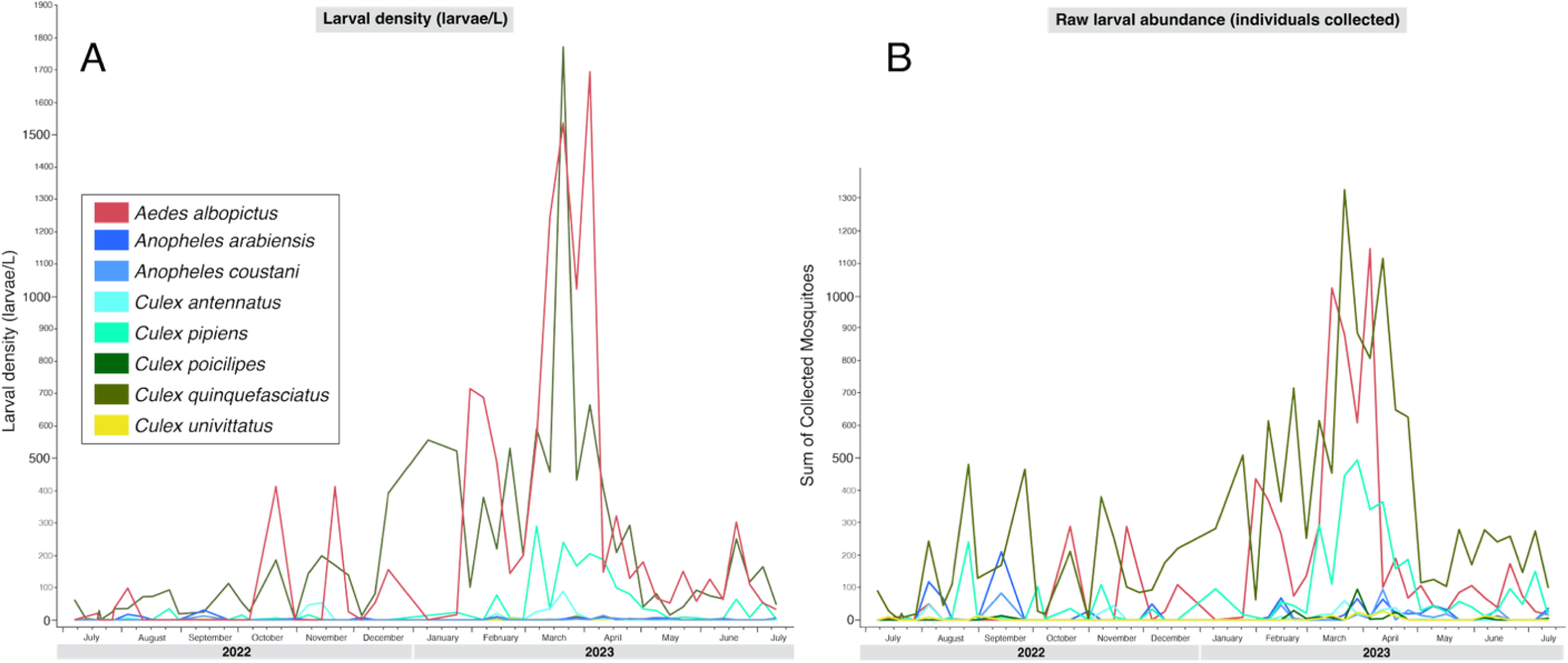
Monthly abundance of the mosquito species in Antananarivo. Larval density (larvae/L; left panel, A) and raw larval abundance (individuals collected per month; right panel, B) are shown as a side-by-side comparison from 2022-2023.

**Figure 4.**
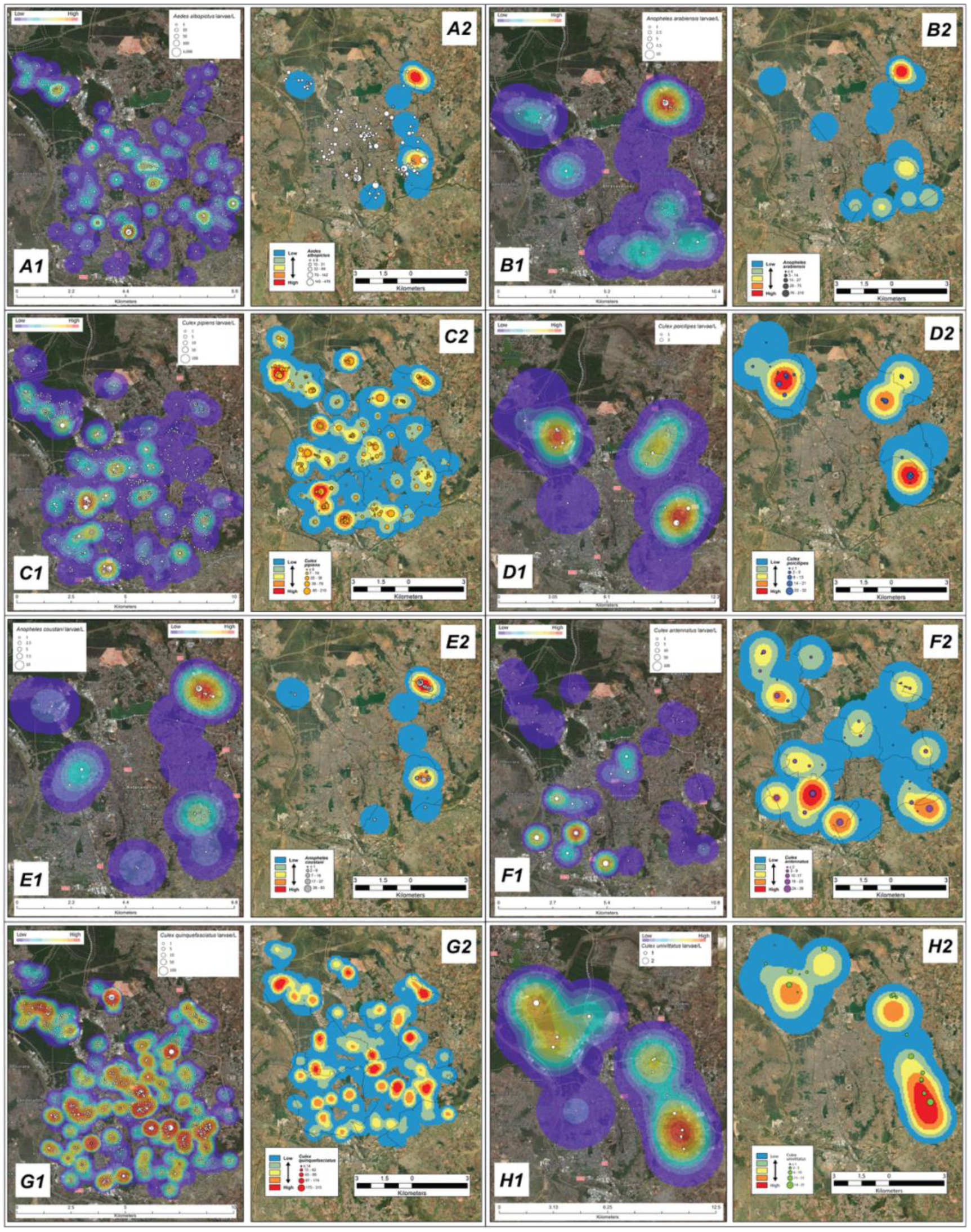
Point and Kernel density distributions of collected mosquitoes by species (A = *Aedes albopictus*, B = *Anopheles arabiensis*, C = *Culex pipiens*, D = *Culex poicilipes*, E = *Anopheles coustani*, F = *Culex antennatus*, G = *Culex quinquefasciatus*, and H = *Culex univittatus*). For direct comparison, larval density (larvae/L) is shown in the left maps (denoted as X1), and raw larval abundance (individuals collected) in the right maps (denoted as X2). Because *Anopheles* occupy large natural habitats, their larval density appears low relative to their raw abundance.

### Monthly fluctuations in vector abundance

The abundance of *Cx. quinquefasciatus* varied from 150 to 1,000 individuals per breeding site during July to January. Populations increased significantly during the rainy season (beginning in February), reaching a peak of 3,800 individuals in March. A slight decrease was observed in April, followed by 1,000 individuals in May and June, and 370 individuals in July 2023. Similar patterns were observed for *Cx. pipiens*, but with lower than 250 individuals for all the months except for March and April, with peaks of 1300 and 1000 individuals, respectively. Low abundance of *Cx. antennatus*, *Cx. poicilipes*, and *Cx. univittatus* were observed for all months, with peaks of approximately 100 individuals recorded in March for the former two species. *Cx. univittatus* peak abundance (50 individuals) occurred in April. The peak abundance of *An. arabiensis* was observed in September, with ∼200 individuals. This abundance remained constant from February through May, ranging between 80 and 120 individuals. Conversely, abundance was low or even zero for the remaining months (June through December, except for August and September). *An. coustani* was marked by two peaks, one in September with 80 individuals and another in April with 128 individuals. This species was always present from February through July, but in low numbers (n < 50 individuals). *An. coustani* was absent from August and October through January. As with the previous species, the abundance of *Ae. albopictus* varied according to the month. Between July and January, the total abundance never exceeded 500 individuals. Conversely, between February and March, the total abundance peaked at 2,800 individuals per month. From April to June, total abundance gradually decreased from 1,500 to <500 individuals (**Figure 3)**. Expressed as larval density, the seasonal signal was similar in timing but was dominated by container-breeding species, with the highest larval densities (larvae/L) of *Ae. albopictus* and *Cx. quinquefasciatus* during the January–March rainy season (Figure 3A), when small artificial containers were most productive per unit volume.

Among the mosquito larvae collected, *Cx. quinquefasciatus* were the most abundant across all of Antananarivo’s arrondissements, especially in arrondissements 2 and 5 (500 and 3300 individuals, respectively). In contrast, the total abundance in arrondissements 1 and 6 was 1500 and <500 individuals, respectively. The median *Ae. albopictus* abundance was 2000 individuals, in arrondissement 2. The number of *Cx. pipiens* is almost the same across all arrondissements, averaging about 500 individuals, except for arrondissement 4, which had a higher abundance (n = 950 individuals). Arrondissements 2 and 5 are also marked by the presence of *An. arabiensis* and *An. coustani* which are almost absent in the other arrondissements. In addition, these arrondissements host the most species (8) compared to the others.

In general, the abundance of mosquitoes collected between the wet season (November to March, n = 14,427 individuals) and the dry season (April to October, n = 13,080 individuals) is not significantly different. The seasonal variation in the abundance of species belonging to the genus *Culex* is not easily distinguishable. In contrast, the abundance of *An. arabiensis*, *An. coustani* and *Ae. albopictus* varies considerably by season. Both *Anopheles* species are abundant during the dry season, while *Ae. albopictus* is abundant year-round; however, *Ae. albopictus* shows higher abundance during the wet season (n = 4,655) compared to the dry season (n = 2,635). A comprehensive summary of the descriptive statistics is found in **Tables S1 & S2**.

### Spatial heterogeneity of vector species

Kernel density distributions of each species display a singular intense concentration of *Ae. albopictus* (**Figure 4A**) and *Cx. antennatus* (**Figure 4F**) in opposing arrondissements. The Kernel density distribution for *Cx. quinquefasciatus* (**Figure 4G**) shows several distinct, intense distributions of collections across all arrondissements.

### Socioenvironmental factors associated with vector species

Results of univariate comparisons of predictor strength for each dependent variable (mosquito species) consistently revealed nine variables with the strongest associations (**Table 1**, **Figure 5**). In particular, the type of larval habitat predictor (e.g., rice field, tire, flower pot, water storage well, bucket, etc.) was found to be the most important covariate in all species analyses except for *An. coustani*. The remaining top predictors (in descending order) for all species were mean imperviousness, mean relative wealth index, mean number of buildings in proximity, human population density, month, mean precipitation, date, and arrondissement. Results from the bootstrap forest model output for all mosquito collections and by each species reveal that the strongest predictions were correlated with increasing the number of collected specimens (**Table S3A**). Bootstrap forest methods applied to all collected mosquitoes (pooled) yielded the strongest-performing model (r^2^ = 0.978, RASE = 3.1). By species, *Cx. quinquefasciatus*, *Cx. pipiens*, and *Cx. poicilipes* were the next three highest-performing models (r^2^ = 0.645, 0.558, and 0.539, respectively). Like the univariate analysis of predictors, the output of bootstrap forest models indicates a similar relationship of each covariate by species (**Table 2)** (full details available in **Table S3B**). Overall, only *An. arabiensis*, *An. coustani, Cx. pipiens*, and *Cx. quinquefasciatus* were significantly spatially autocorrelated, with a clustered pattern (**Table 3**).

**Figure 5.**
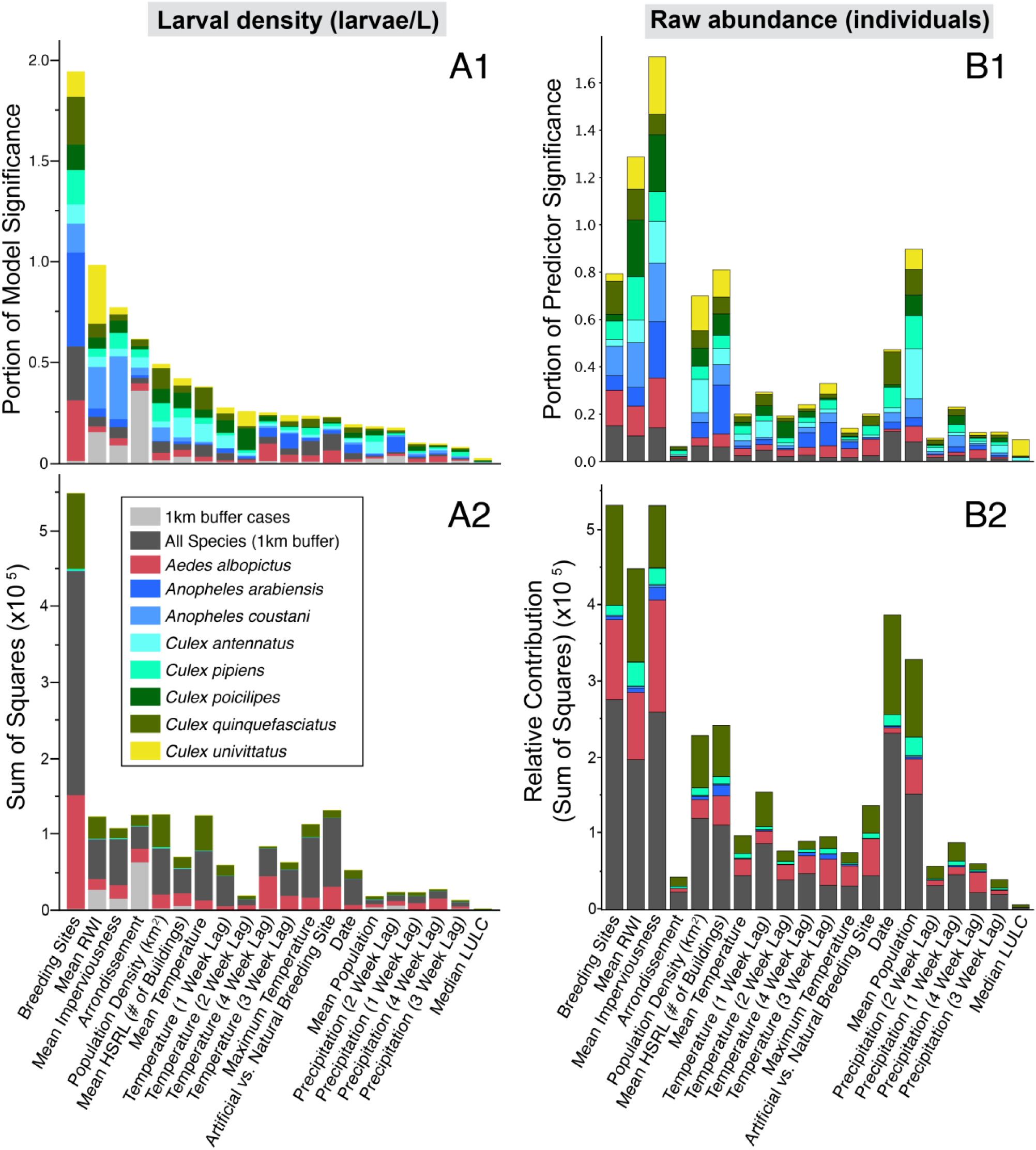
Comparison of univariate predictor strength for each mosquito species (stacked accumulations), assessed by model significance portion (top) and sum of squares (bottom). Larval density (larvae/L; A) and raw larval abundance (individuals; B) analyses are shown side by side; larval-habitat type is the leading predictor of larval density, whereas urban covariates (mean imperviousness, relative wealth index) rank higher for raw abundance.

**Table 1.** Univariate comparisons of predictor strength by dependent variable. Overall rankings are sorted by descending average rank (importance) across all individual species (dependent variable) comparisons.

| Species | Top 3 Predictors (by descending rank) |  | Contribution | Portion |
| --- | --- | --- | --- | --- |
| All Human Cases<br>(1km buffer) | #1 | Arrondissement | 59876.8 | 0.3444 |
|  | #2 | Mean RWI | 29274.9 | 0.1684 |
|  | #3 | Mean Imperviousness | 14863.9 | 0.0855 |
| All Species | #1 | Larval Habitats | 392430 | 0.3482 |
|  | #2 | Population Density (km<sup>2</sup>) | 74111 | 0.0658 |
|  | #3 | Mean Temperature | 61436 | 0.0545 |
| *Aedes albopictus* | #1 | Larval Habitats | 135510 | 0.2808 |
|  | #2 | Temperature Lag 4 weeks | 27727 | 0.0574 |
|  | #3 | Artificial vs Natural Larval habitats | 26658 | 0.0552 |
| *Anopheles arabiensis* | #1 | Larval Habitats | 176.099 | 0.4751 |
|  | #2 | Temperature Lag 3 weeks | 29.460 | 0.0795 |
|  | #3 | Date | 21.258 | 0.0573 |
| *Anopheles coustani* | #1 | Mean Imperviousness | 18.3335 | 0.2801 |
|  | #2 | Mean RWI | 15.4527 | 0.2361 |
|  | #3 | Larval Habitats | 9.3434 | 0.1427 |
| *Culex antennatus* | #1 | Larval Habitats | 257.053 | 0.1056 |
|  | #2 | Children (< 5 years old) | 210.037 | 0.0863 |
|  | #3 | Mean Temperature | 201.354 | 0.0827 |
| *Culex pipiens* | #1 | Larval Habitats | 2445.97 | 0.1696 |
|  | #2 | Population Density (km<sup>2</sup>) | 1174.33 | 0.0814 |
|  | #3 | Mean Imperviousness | 1109.61 | 0.0769 |
| *Culex poicilipes* | #1 | Larval Habitats | 2.84559 | 0.1550 |
|  | #2 | Temperature Lag 2 weeks | 2.11848 | 0.1154 |
|  | #3 | Population Density (km<sup>2</sup>) | 1.92582 | 0.1049 |
| *Culex quinquefasciatus* | #1 | Mean Imperviousness | 104536 | 0.2451 |
|  | #2 | Mean Temperature | 48090 | 0.1127 |
|  | #3 | Population Density (km<sup>2</sup>) | 46006 | 0.1079 |
| *Culex univittatus* | #1 | Mean RWI | 2.80101 | 0.3075 |
|  | #2 | Larval Habitats | 1.26096 | 0.1384 |
|  | #3 | Temperature Lag 2 weeks | 0.53557 | 0.0588 |
| Average Rank | #1 | Larval Habitats | 1.38 |  |
|  | #2 | Mean RWI | 4.75 |  |
|  | #3 | Mean Imperviousness | 6.25 |  |

**Table 2.** Summary of Bootstrap Forest model output and breakdown of the top 3 highest-performing covariates by mosquito species (dependent variable).

| Species | Top 3 Predictors (by descending rank) |  | Number of Splits | Contribution | Portion |
| --- | --- | --- | --- | --- | --- |
| All Human Cases<br>(1 km buffer) | #1 | Arrondissement | 342 | 61946.37 | 0.3541 |
|  | #2 | Mean RWI | 1275 | 25377.23 | 0.1451 |
|  | #3 | Mean Imperviousness | 993 | 15471.09 | 0.0884 |
| All species | #1 | Larval Habitats | 1261 | 286056.69 | 0.2585 |
|  | #2 | Artificial vs Natural Larval Habitats | 133 | 95352.15 | 0.0862 |
|  | #3 | Population Density (km<sup>2</sup>) | 583 | 94143.63 | 0.0851 |
| *Aedes albopictus* | #1 | Larval Habitats | 774 | 154100.46 | 0.3237 |
|  | #2 | Temperature Lag 4 Weeks | 171 | 34123.39 | 0.0717 |
|  | #3 | Artificial vs Natural Larval Habitats | 129 | 29078.27 | 0.0611 |
| *Anopheles arabiensis* | #1 | Larval Habitats | 250 | 227.20 | 0.5552 |
|  | #2 | Temperature Lag 3 Weeks | 83 | 36.68 | 0.0896 |
|  | #3 | Date | 74 | 25.11 | 0.0614 |
| *Anopheles coustani* | #1 | Mean Imperviousness | 310 | 20.72 | 0.3027 |
|  | #2 | Larval Habitats | 180 | 10.30 | 0.1505 |
|  | #3 | Mean RWI | 130 | 10.12 | 0.1478 |
| *Culex antennatus* | #1 | Temperature Lag 1 Week | 57 | 295.48 | 0.1224 |
|  | #2 | Larval Habitats | 170 | 230.35 | 0.0954 |
|  | #3 | Mean HSRL (# of Buildings) | 113 | 156.53 | 0.0648 |
| *Culex pipiens* | #1 | Larval Habitats | 902 | 2341.71 | 0.1679 |
|  | #2 | Population Density (km<sup>2</sup>) | 390 | 1085.61 | 0.0778 |
|  | #3 | Mean Imperviousness | 643 | 984.72 | 0.0706 |
| *Culex poicilipes* | #1 | Larval Habitats | 89 | 1.98 | 0.1186 |
|  | #2 | Temperature Lag 2 Weeks | 52 | 1.58 | 0.0948 |
|  | #3 | Mean RWI | 107 | 1.50 | 0.0897 |
| *Culex quinquefasciatus* | #1 | Larval Habitats | 1217 | 104457.02 | 0.2571 |
|  | #2 | Population Density (km<sup>2</sup>) | 481 | 37429.13 | 0.0921 |
|  | #3 | Mean Temperature | 248 | 28032.99 | 0.0690 |
| *Culex univittatus* | #1 | Mean RWI | 135 | 2.52 | 0.2854 |
|  | #2 | Larval Habitats | 88 | 1.32 | 0.1492 |
|  | #3 | Temperature Lag 2 Weeks | 33 | 0.65 | 0.0735 |

**Table 3.**
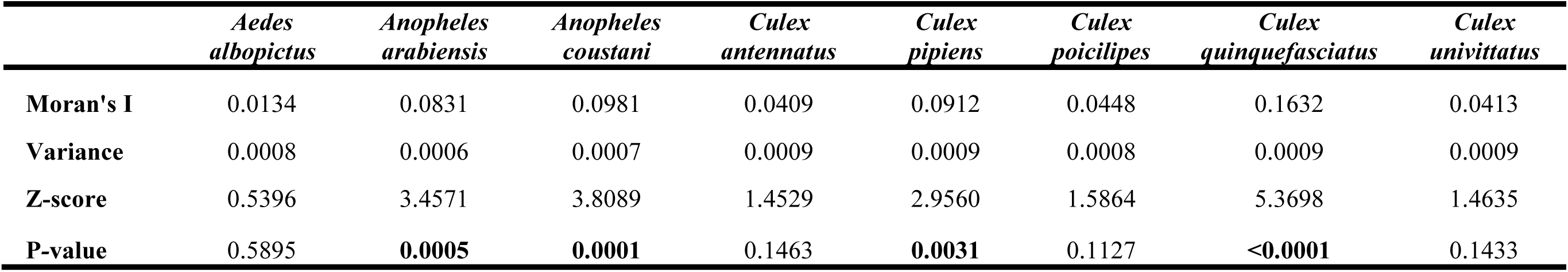
Assessment of spatial autocorrelation for all collected mosquitoes, by species, in Antananarivo, Madagascar, from July 2022 – July 2023. Moran’s I can be interpreted as follows: values closer to +1 indicate clustering; values closer to -1 indicate dispersion; values close to 0 indicate random spatial relationships (no autocorrelation). In Antananarivo, *Aedes albopictus, Culex antennatus,* and *Culex quinquefasciatus* demonstrate a statistically significant clustering relationship.

## Discussion

Urbanization has the potential to facilitate transmission of MBDs such as dengue, chikungunya, (WNV) virus, and malaria through expanded human population density, increased abundance and density of vector larval habitats, larval density, and microclimate impacts. The increase in the urban form of malaria has been recognized by the WHO as a threat to eliminating the parasite. These threats are exacerbated by invasive malaria vectors that thrive in urban areas, in particular *An. stephensi* (WHO Urban Malaria Framework, 2022).

Antananarivo is the capital and largest urban center on the island of Madagascar, and our year-long urban mosquito larval surveillance study was conducted across urban and peri-urban gradients throughout the city. Molecular methods for identifying *Anopheles* species were used to validate the potential for citizen science- and artificial intelligence-based larval surveillance in the future (Carney *et al*., 2022, 2023), as well as to determine trends in mosquito vector dynamics to better understand the seasonal risk of MBDs and to develop data-driven vector control strategies. The findings presented here identify dynamics in both spatial and temporal preferences, highlighting key characteristics of suitable habitats by mosquito vector species. All vectors increased in abundance during the rainy season, especially from January to March; *An. arabiensis*, one of the primary malaria vectors in Madagascar, showed two peaks in abundance: one during the dry season (July-October) and a second during the rainy season; however, the species persisted year-round, and no significant seasonal differences were found. This highlights the risk of malaria and MBD year-round in an urban setting where larval habitats persist. Unlike reports of typical malaria vectors in Madagascar, *An. arabiensis* in Antananarivo was also abundant in other habitats, particularly brick pits, wells, and water basins. The use of these habitats may help explain dry-season persistence and pose an urban malaria threat.

Distinguishing larval density from raw abundance also has direct operational relevance for vector control. Locations where *Anopheles* and *Aedes* larval habitats overlap—evident in the spatial distributions in Figure 4 and concentrated within specific arrondissements—represent priority areas for integrated vector management, where a single set of interventions can simultaneously reduce malaria and arboviral risk (WHO, 2026). Because larval density is highest in small, manageable container habitats, these results pinpoint the exact localities and arrondissements where larval source management (LSM), including container reduction and targeted larviciding, is most likely to be cost-effective, consistent with current WHO operational guidance (WHO, 2026). Finally, the temporal resolution of this surveillance underscores the value of real-time larval monitoring, enabling control measures to be implemented and evaluated during periods and in areas of peak transmission risk to protect human health.

This study shows that the landscape of Antananarivo’s urban area is highly heterogeneous, comprising a multitude of larval habitats: 55% are of artificial origin, and 45% are natural. In three examples, positive larval habitats of *Cx. antennatus*, *Cx. pipiens*, and *Cx. quinquefasciatus* were frequently collected throughout all arrondissements. Conversely, *Ae. albopictus*, *An. arabiensis*, *An. coustani, Cx. poicilipes,* and *Cx. univittatus* were collected in markedly fewer locations, albeit with one or two areas of high density. The larval habitats observed in the six arrondissements of the city are very diverse but can be grouped into 21 types: bamboo, brick pit, coconut waste, cut plastic canister, drainage channel, flower pot, footprint, metal container, plastic bucket, plastic bowl barrel, pond, puddle, rice field, small plastic container, tire, tree tank, used bottle, water basin, watercress and water well. Most of these larval habitats are associated with human activities in urban areas, such as agriculture, construction, and aquatic waste (WHO, 1982; Minakawa *et al*., 1999; Li *et al*., 2014; Mayi *et al*., 2020). All these activities generate stagnant water, which is favorable to the development of mosquito larvae. Karuitha and colleagues suggested that human behavior and socioeconomic parameters play an important role in the generation of larval habitats (Woolhouse & Gowtage, 2005; Karuitha *et al*., 2019).

Eight species across three genera were identified, suggesting relatively high mosquito diversity, consistent with the heterogeneity of urban larval habitats. Similar observations have been reported in other studies, which also documented high mosquito diversity in urban areas (Egbuche *et al*., 2016; Onyekachi *et al*., 2018; Amusan & Ogbogu, 2020). Herein, *Culex* larvae dominate in terms of abundance and diversity, with five species compared to *Aedes* (one species) and *Anopheles* (one species). Previous studies have confirmed this observation by reporting higher *Culex* abundance in urban and peri-urban zones during larval surveillance (Karuitha *et al*., 2019; Amusan & Ogbogu, 2020). *Cx. quinquefasciatus* (53%, n = 14758) and *Cx. pipiens* (14%, n = 3746) represent the majority of species in this genus. The high abundance of the latter in the urban environment can be explained by their ability to reproduce in a wide range of larval habitats that are characterized by polluted waters with unpleasant odors, as mentioned by some authors (Urbanelli *et al*., 1995; Mafiana *et al*., 1998; Noori *et al*., 2015; Okwa *et al*., 2018).

Indeed, these larvae serve as bioindicators of water quality, as their presence can be associated with polluted habitats. In this study, *Cx. quinquefasciatus* is present in almost all types of larval habitats encountered throughout all of Antananarivo, although a particular preference for watercress and puddle habitats has been observed. This suggests that the species is highly adaptable and can colonize all existing larval habitats in urban environments (Karuitha *et al*., 2019). The transformation of rice fields into watercress in Antananarivo (Aubry *et al*., 2011) is suggested as favoring the abundance of *Cx. quinquefasciatus*. The highest prevalence of this species in the same city was also reported by other authors (Fontenille, 1989; Ravoahangimalala *et al*., 2008; and Raharimalala *et al*., 2012). This confirms this species’ status as an urban species in tropical regions, South Asia, and the Mascarene archipelago (WHO, 1988; Tantely *et al*., 2010). In some countries, such as Kenya and Nigeria, the urban environment is largely dominated by *Cx. pipiens* instead of *Cx. quinquefasciatus* (Karuitha *et al*., 2019; Amusan & Ogbogu, 2020). *Cx. univittatus*, *Cx. poicilipes*, and *Cx. antennatus* are a minority, with a proportion of about 1%, and have been found only in some larval habitats, such as rice fields and brick pits. This implies that the larvae of these species are closely associated with rice cultivation, which occurs mainly in the peri-urban area (Robert *et al*., 2002). Their larval habitats are characterized by less polluted waters than those of the two species mentioned above. The presence of *Cx. antennatus* in tires and cut plastic canisters probably indicates that it has adapted over time to colonize domestic larval habitats near dwellings, but a more in-depth study is needed to confirm this hypothesis.

The results showed that *Cx. pipiens* and *Cx. quinquefasciatus* present a high larval density during the wet season. The relevant explanation for this observation is the arrival of rains, which fill all potential larval habitats and significantly increase the number of larvae in the study area. During the dry season, the juvenile stages of these two species are still present but at low abundance. It is assumed that only the permanent larval habitats remain in the city’s different regions during this season, and that these habitats sustain mosquito larval populations by serving as reservoirs. The same findings were recorded for *Cx. univittatus*, *Cx. poicilipes*, and *Cx. antennatus*, which presents more larvae during the wet season. These observations are the opposite of those reported by other studies that found that mosquito abundance was greater in the dry season than in the wet season (Okwa *et al*., 2018; Amusan & Ogbogu, 2020); they explain that many larvae and potential larval habitats are washed away by rain runoff, causing a reduction in the mosquito population during the wet season.

The high prevalence of *Culex* species, in particular *Cx. quinquefasciatus* in urban environments undoubtedly constitutes a significant source of nuisance to residents through bites inflicted during the night and their anthropophilic behavior (Fontenille, 1989). This species is known as the main vector of *Bancroftian filariasis* worldwide (Mwakitalu *et al*., 2013; Derua *et al*., 2017) and can also transmit the Rift Valley Fever virus (RVFV) (Ndiaye *et al*., 2016), WNV, and many other parasites and viruses. Thus, the existence of *Cx. quinquefasciatus* throughout the year in Antananarivo constitutes a high risk of transmission, especially in March, which corresponds to its peak abundance. *Cx. pipiens* has been reported as a vector of WNV (Kwan *et al*., 2010) and RVFV (Moutailler *et al*., 2008; Brustolin *et al*., 2017). Considered to be a zoophilic species, Tantely et al. (2016) suggested that *Cx. pipiens* is not involved in disease transmission in Madagascar (Tantely *et al*., 2016), but its vector capacity should be investigated, especially in urban areas such as Antananarivo. The low abundance of larvae of *Cx univittatus*, *Cx poicilipes*, and *Cx antennatus* in the study area is not negligible in terms of medical importance, as they have been reported as vectors of pathogens affecting humans, namely *Wuchereria bancrofti* (Brunches, 1969), RVFV (Ratovonjato *et al*., 2010; Tantely *et al*., 2013), WNV (Fontenille, 1989), and other viruses (Khan *et al*., 2017; Mavridis *et al*., 2018; Patsoula *et al*., 2020).

The genus *Aedes*, represented only by *Ae. albopictus*, is the second most abundant species in Antananarivo, with a 26% (n = 7,290) prevalence, of which approximately 88% (n = 6,420) were collected from artificial larval habitats near human dwellings. This clearly shows that the abundance of this mosquito is closely linked to the development of domestic larval habitats, namely, used or abandoned containers such as discarded bottles that account for 64% of household waste (Raharinjanahary *et al*., 2011). This is why *Ae. albopictus* has long been considered an urban species (WHO, 1988; Tantely *et al*., 2010). These observations confirm those of previous studies that reported a similarity of results concerning the dominance of this mosquito in urban environments, particularly around human habitats (Senthamarai & Jebanesan, 2016; Amusan & Ogbogul, 2020). The larval habitats of *Ae. albopictus* are generally characterized by fairly clean, less polluted waters (Karuitha *et al*., 2019) and are rich in plant debris such as tires, flower pots, and bamboo. *Ae. albopictus*, known as a highly invasive species (Global Invasive Species Database; Bradshaw *et al*., 2016), can colonize various larval habitats, even small puddles of water in a container. This is the case for tree tanks, plastic buckets, and cut plastic canisters observed in some localities of Antananarivo. Studies have reported that several factors, such as temperature, relative humidity, and water quality, affect the survival and proper development of this species in larval habitats (Chen, Lee *et al*., 2009; Dejene *et al*., 2015). The adaptability of *Ae. albopictus* in different aquatic habitats in the urban area confirms its strong ecological plasticity (Boubidi, 2016). The rainy season is the period during which the highest abundance of this mosquito’s larvae was recorded in Antananarivo, peaking in March, as with the other species mentioned above. However, it remains at low abundance during the dry season thanks to the ability of its eggs to resist desiccation and enter diapause (Hawley *et al*., 1987; Hanson & Craig, 1994; Lacour, 2016). In the presence of a small amount of water, the eggs resume their development into larvae. Indeed, this allows *Ae. albopictus* to ensure its sustainability in the face of environmental and climatic variations. This study highlights that urban areas offer a variety of larval habitats suitable for the development of *Ae. albopictus* larvae. This is essential information because the adult female of this species is known from laboratory experiments to be competent to transmit at least 26 different viruses, such as dengue and Chikungunya (Delatte *et al*., 2008; Ratsitorahina *et al*., 2008; Bagny *et al*., 2009) and Zika (Grard *et al*., 2007; Paupy, 2010; Wong *et al*., 2013; Kraemer *et al*., 2017). Apart from its strong anthropophagy in urban environments (Richards *et al*., 2006; Valerio *et al*., 2010; Kamgang *et al*., 2012), *Aedes* can take multiple blood meals (Hawley, 1988; Ponlanwat & Harrington, 2005) and exhibit aggressive biting behavior during the day (Delatte *et al*., 2010). For these reasons, the risk of pathogen transmission to residents is higher in urban areas, while control will be difficult in the event of an epidemic due to the high density of the urban population and the substantial diversity of larval habitats.

Two species of *Anopheles* were identified in this study, *An. coustani* and *An. arabiensis* (confirmed by molecular analysis). They were collected from eight different larval habitats, including brick pit, footprint, pond, puddle, rice field, water basin, watercress, and water well. Among these larval habitats, the two species showed a high abundance in rice fields and brick pits. This assumes that the latter are the preferred larval habitats of these mosquitoes in urban areas. A significant abundance of *An. coustani* in rice fields in the peri-urban areas of Antananarivo has also been mentioned by previous studies (Fontenille, 1989; Tantely *et al*., 2013). Apart from the larval habitats reported in this study, the larvae of this species can also be observed in various other habitats such as cattle hoof prints (Doucet, 1949), ponds, swamps (Grejbine, 1953), rivers, streams, canals (Doucet, 1951), lakes, rock holes, flushing holes (Le Goff, 1951), and brackish water pools (Grejbine, 1966) in many regions of Madagascar. *An. coustani*, known as a zoo-anthropophilic species, can transmit infectious agents of malaria, including *Plasmodium falciparum* and *P. vivax* (Nepomichene *et al*., 2015; Goupeyou - Youmsi *et al*., 2020), *Wuchereria bancrofti* (Brunches, 1969), and several viruses such as RVFV (Ratovonjato *et al*., 2010) and WNV (Maquart *et al*., 2016) in Madagascar. According to our results, the main breeding site of *An. arabiensis* is a brick pit. This type of larval habitat is characterized by a shady, undisturbed environment. Tantely et al. reported that larvae of this species develop in freshwater larval habitats (Tantely *et al*., 2016).

A dominance of *An. arabiensis* compared to *An. coustani* was found in this study. Similar observations have emphasized the predominance of this species in some localities of the city of Antananarivo (Fontenille *et al*., 1990; Léong Pock Tsy *et al*., 2003; Rabarijaona *et al*., 2006; Ravoahangimalala *et al*., 2008). This suggests that over time, the *An. gambiae* complex is still represented by the single species *An. arabiensis* in the central highlands of Madagascar, especially in urban and peri-urban areas. It has been reported that *An. arabiensis* plays an important role in the transmission of malaria in Madagascar (Léong Pock Tsy *et al*., 2003; Ravoahangimalala *et al*., 2008; Ratovonjato *et al*., 2014), although it exhibits a low degree of anthropophilic behavior (Ralisoa, 1996). Both Anopheles *species* are significantly abundant during the dry season, and their monthly variation shows two peaks in September and April. It is assumed that this is due to the availability of rice fields and brick pits, which are numerous during the dry season due to agricultural and construction activities in the study area. On the other hand, the washing of larvae by heavy rains in the larval habitats largely explains the low abundance of *An. arabiensis* and *An. coustani* during the wet season. Unlike the genera *Aedes* and *Culex*, *Anopheles* larvae are much more sensitive to habitat variations, as their development can be affected by factors such as water temperature, larval density, and nutrient availability (Spitzen & Takken, 2005; Mamai *et al*., 2018). Thus, they require larval habitats with appropriate conditions.

The intensive, year-long collection campaign generated tens of thousands of mosquito specimens across all arrondissements. However, apparent mosquito abundance was heavily skewed by species, as *Cx. quinquefasciatus, Cx. pipiens,* and *Ae. albopictus* accounted for 93.0% of all collections. As expected, species with far fewer collected individuals resulted in poorer model fit predictions. This trend does not apply to *Cx. poicilipes*, however. Despite collecting only 200 individuals, the Bootstrap Forest model had the third-highest performance across all species. However, this is likely an artifact of collections by habitat type (93.5% were collected in rice fields) and should be further investigated to determine whether other factors contributed to the higher-than-average model performance. Future analyses by similar studies should be conducted, incorporating model-fit parameters specific to the biological and behavioral preferences of each species of interest.

These results indicate species-specific preferences for breeding habitat type, seasonality, and other environmental conditions. Additionally, positive larval habitats throughout the study area suggest spatially unique “hot spots” that generate large collections for some species. These sites should be investigated more closely to identify potential microclimatic and other unique habitat conditions that may be conducive to reproduction. These findings have the potential to provide valuable “ground-truth” validation for refining each species’ model fit while simultaneously implementing highly efficient, targeted fine-scale interventions for vector control.

## Acknowledgments

We would like to thank the weather stations in Antananarivo for providing the temperature and precipitation data used in this study. We also thank the International Associated Laboratory that allowed us to rear mosquitoes in their insectary.

## Disclaimer

The findings and conclusions expressed herein are those of the authors and do not necessarily represent the official position of the Centers for Disease Control and Prevention (CDC).

